# Genetic diversity of wild and cultivated *Coffea* canephora in northeastern DR Congo and the implications for conservation

**DOI:** 10.1101/2021.08.09.455630

**Authors:** Samuel Vanden Abeele, Steven B. Janssens, Justin Asimonyio Anio, Yves Bawin, Jonas Depecker, Bienfait Kambale, Ithé Mwanga Mwanga, Tshimi Ebele, Salvator Ntore, Piet Stoffelen, Filip Vandelook

## Abstract

**Premise:** Many cultivated coffee varieties descend from *Coffea canephora*, commonly known as Robusta coffee. The Congo Basin has a century long history of Robusta coffee cultivation and breeding, and is hypothesized to be the region of origin of many of the cultivated Robusta varieties. Since little is known about the genetic composition of *C. canephora* in this region, we assessed the genetic diversity of wild and cultivated *C. canephora* shrubs in the Democratic Republic of the Congo.

**Methods:** Using 18 microsatellite markers, we studied the genetic composition of wild and backyard-grown *C. canephora* shrubs in the Tshopo and Ituri provinces, and from the INERA Yangambi Coffee Collection. We assessed genetic clustering patterns, genetic diversity, and genetic differentiation between populations.

**Key results:** Genetic differentiation was relatively strong between wild and cultivated *C. canephora* shrubs, and both gene pools harbored multiple unique alleles. Strong genetic differentiation was also observed between wild populations. The level of genetic diversity in wild populations was similar to that of the INERA Yangambi Coffee Collection, but local wild genotypes were mostly missing from that collection. Shrubs grown in the backyards were genetically similar to the breeding material from INERA Yangambi.

**Conclusions:** Most *C. canephora* that is grown in local backyards originated from INERA breeding programs, while a few shrubs were obtained directly from surrounding forests. The INERA Yangambi Coffee Collection could benefit from an enrichment with local wild genotypes, to increase the genetic resources available for breeding purposes, as well as to support ex situ conservation.

## INTRODUCTION

Coffee is one of the most valuable crops in the world and is the second-most exported product of developing countries (Pendergast, 2009). Most cultivated coffee varieties descend from two wild species: *Coffea arabica* L. (Arabica coffee) and *C. canephora* Pierre ex A.Froehner (Robusta coffee), of which the latter represents close to 44 % of the global coffee production (data provided by ICO, statistical service). Whereas Arabica coffee was introduced by Arabian merchants in Yemen for cultivation c. 1000 years ago (Smith, 1985), the commercial cultivation of Robusta coffee is less than 150 years old. In the late 19th century, reports on the local cultivation of *Coffea canephora* were made from Gabon, Angola and Uganda (pers. observ. by PS on herbarium label and Chevalier, 1929). However, widespread colonial cultivation of Robusta coffee started only in the early twentieth century. The introduction and promotion of ‘*Coffea robusta’* as a robust coffee species by the Belgian horticulturist Linden in 1900 is probably key for the success of Robusta coffee, as the commercial name is suggesting. Linden’s introduction was done using seeds of wild plants from the Sankuru province in the Democratic Republic of the Congo (DR Congo). This material was sent to Java, where it was crossed with other Robusta lineages i.a. from Lower Congo and Uganda. After the arrival of ‘Robusta coffee’ in Java in the early 20th century, Java developed itself to become an important breeding and distribution center of Robusta coffee (Ferrão et al., 2019). Before the European colonization of Africa, *Coffea canephora* was only grown locally (Jaroget and Descroix, 2002), mainly in the northeastern and southwestern part of its natural distribution area.

In the early 1900’s, the first Robusta coffee research and breeding stations were also installed in Central Africa, e.g. the Botanical Garden in Eala. In DR Congo, the INEAC (Institut National pour l’Etude Agronomique du Congo Belge) was created in 1933, to develop a program for scientific research focused on agriculture and forestry, with a network of research stations throughout the country (Jaroget and Descroix, 2002). Yangambi (Tshopo province, northeastern DR Congo) became the principal research station of the INEAC, both in general and for Robusta coffee (Leplae, 1936). In the years following the second World War, DR Congo and Uganda took over Java’s role as principal research and breeding centers for *C. canephora* (Jaroget and Descroix, 2002). From there, plants and seeds were distributed to other regions. In this context, INEAC Yangambi also assembled a large Robusta coffee gene bank. In 1962, two years after the Independence of Congo, INEAC changed to become the INERA (Institut National des Etudes et Recherches Agronomiques). After this change, the research and breeding activities at INERA Yangambi were gradually reduced and during the last decades, many accessions of the gene bank were lost. In 2016, the Robusta Coffee Collection of the INERA Yangambi held 94 different genetic lines, of which seven were elite breeding lines (6 Lula & 1 Java line; pers. observ. FV & PS). Currently, important Robusta research centers are situated in Brazil, Vietnam, Uganda and India. Valuable Robusta genetic resources, including material originating from the INEAC station in Yangambi, are held in collections in Cameroon, Ivory Coast, India and Madagascar (Cubry et al., 2013; Bramel et al., 2017).

In contrast to *C. arabica* (Aerts et al., 2013), it can be expected that vast amounts of untouched wild genetic diversity of *C. canephora* still exist across its wide distribution range. Wild *C. canephora* occurs in the rainforests of West and Central Africa, from Guinea to Uganda, occupying the largest distribution area among all *Coffea* species (Noirot et al., 2016). A recent molecular study demonstrated the presence of eight clearly delineated genetic clusters in wild *Coffea canephora* populations (Merot-L’anthoene et al., 2019). One of these genetic clusters roughly encompasses the northeastern part of the Congo Basin, including the Yangambi area (Tshopo province). The cluster covers a large area Northeast to the Congo River and the city of Kisangani. Vegetation in this area is characterized by both old-growth and intervened forests (Gilson, 1956) in which wild *C. canephora* populations are present as understory shrubs, often sympatric with *Coffea liberica* and *Coffea dactylifera* (pers. obs.). *Coffea canephora* shrubs typically grow at low density in small, disconnected populations (Musoli et al., 2009). Information on population genetic diversity and structure of wild *C. canephora* is scarce and, as far as we know, not available for the DR Congo.

The Congo Basin is hypothesized to be the region of origin of many cultivated Robusta coffee genotypes (Dulloo et al., 1998; Cubry et al., 2013). Consequently, populations of *C. canephora* native to this region contain a valuable part of the wild gene pool, but the extent of this genetic reservoir remains unknown. In Uganda, comparison of cultivated Robusta accessions with wild *C. canephora* populations indicated a significantly higher genetic diversity among cultivated accessions than in wild populations (Musoli et al., 2009). A recent study has shown that cultivated accessions in Uganda are genetically very similar to wild populations from southwestern Uganda (Kiwuka et al., 2021), suggesting a common genetic origin, recent introduction in cultivation and limited breeding. While in Uganda *Coffea canephora* is mainly cultivated in plantations, this is not the case in the Congolese Tshopo and Ituri provinces. Although *Coffea canephora* plantations could be found throughout the Tshopo and Ituri provinces in the twentieth century, these plantations have disappeared over the last decades. Currently, *C. canephora* shrubs are mostly grown in small-scale backyard garden systems consisting of only a few shrubs for domestic use.

Although the DR Congo potentially harbors an enormous reservoir of genetic diversity of *C. canephora*, and has a century long history of breeding and cultivation, virtually nothing is known about the genetic composition of both wild and cultivated *C. canephora* in this region. Therefore, we present for the first time a study focused on the genetic diversity of *C. canephora* in the DR Congo, in which we apply population genetics methods on wild and backyard-grown shrubs in the Tshopo and Ituri provinces and on the INERA Yangambi Coffee Collection. The following questions will be addressed: (i) Are backyard-grown coffee shrubs genetically different from nearby growing wild shrubs? (ii) How does the genetic diversity compare between cultivated and wild shrubs? (iii) How much (local) genetic diversity is preserved in the Coffee Collection of the INERA Yangambi? (iv) What is the level of genetic differentiation among wild *Coffea canephora* populations? (v) What are the implications for the conservation of *C. canephora* genetic resources in the DR Congo?

## MATERIALS AND METHODS

### Taxon sampling and DNA extraction

Leaf samples of wild and cultivated *Coffea canephora* shrubs were collected at multiple localities in the DR Congo (Figure 1). Natural populations were sampled in the Yangambi and Yoko reserves (both in the Tshopo province), and in Epulu and Djugu (both in the Ituri province). Cultivated specimens were collected from backyards in Yangambi, Kisangani (both in the Tshopo province) and Epulu (Ituri province), very often fairly close to the wild shrubs. The INERA Yangambi Coffee Collection was sampled exhaustively (45 samples). In total, 195 leaf samples were collected (Appendix 1) and dried with silica-gel for molecular analyses. Genomic DNA was isolated using a cetyltrimethylammonium bromide (CTAB) protocol (Doyle and Doyle, 1990) with an additional sorbitol washing step (Janssens et al., 2006).

**Figure 1.**
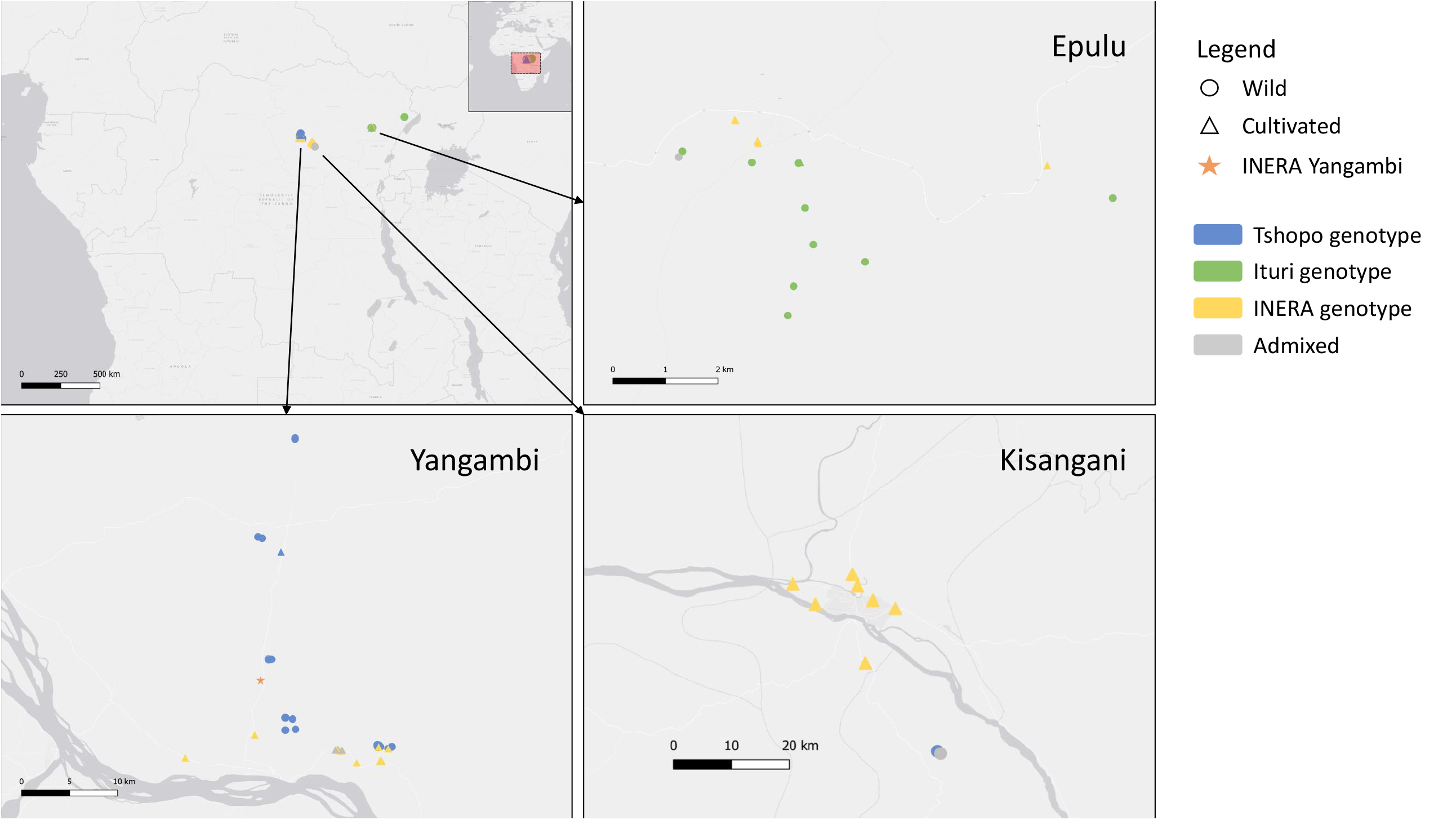
Sampling locations of wild and cultivated *Coffea canephora* (Robusta coffee) in northeastern Democratic Republic of the Congo. The map was made in QGIS 3.4 (QGIS.org, 2021) using the World Light Gray Base layer (esri, 2016).

### Microsatellite primer selection and genotyping

Microsatellite loci amplification was done using 18 primer pairs previously used on wild and cultivated *C. canephora* samples from Uganda (Kiwuka et al., 2021). To reduce the cost of primers, multiplex PCRs were done using an M13-like labelling protocol as described by Schuelke (2000). Therefore, a unique Q-tail sequence (i.e. Q1 after Schuelke (2000), Q2, Q3, or Q4 after Culley et al. (2008)) was added to the 5’ end of the original reverse primers (Appendix S1; see Supplemental Data with this article). The PCR mix (final volume of 15.75 µL) consisted of: 7.5 µL Type-it Multiplex PCR Master Mix (QIAGEN), 3 µL Q solution (5X), 0.3 µL unlabeled forward primer (10 µM), 0.1 µL Q-tailed reverse primer (10 µM), 0.3 µL of a primer (10 µM) composed of the same universal Q1-Q4 sequence with a fluorescent dye attached to the 5’ (6-FAM, NED, VIC and PET, respectively), 1 µL DNA extract, and H_2_O. Multiplex PCR conditions were as follows: initial denaturation at 95 °C (3 min); 25 cycles of denaturation at 95 °C (30 s), annealing at 57 °C (45 s), elongation at 72 °C (1 min); 10 cycles of denaturation at 95 °C (30 s), annealing at 53 °C (45 s), elongation at 72 °C (60 s), and a final extension step at 72 °C (10 min).

Genotyping was done on an ABI 3730 DNA Analyzer (Applied Biosystems) with 1,5 µL PCR product, 12 µL Hi-Di Formamide (Applied Biosystems) and 0.3 µL MapMarker 500 labelled with DY-632 (Eurogentec). Allele calling and locus bin setting was done using the Microsatellite Plugin 1.4.6 in Geneious 9.1.6 (Kearse et al., 2012).

### Genetic population structure and admixture

We used the Bayesian clustering algorithm implemented in the STRUCTURE software v. 2.3.4 (Pritchard et al., 2000) to infer population structure and to assess levels of admixture among cultivated and wild *C. canephora* plants collected in northeastern DR Congo. The following parameters were used: burn-in period and number of MCMC replicates after burn-in both set at 100000, admixture model, independent allele frequency model, maximum number of clusters set between *K* = 1 and *K* = 10, and 10 iterations for each *K*. As recommended by Wang (2017), an alternative ancestry prior α was used, which improves individual assignments and inference of the number of clusters *K*, even if sampling is highly unbalanced (Wang, 2017). This is especially useful in our case, since few samples from Djugu and Yoko were included, and the INERA Yangambi Coffee Collection contains rare specimens from underrepresented localities (Appendix 1). Therefore, the ancestry prior α for each cluster was assumed to be distinct and α was set to an initial value of 0.25 (equals 1/*K*, with *K* = 4 based on preliminary clustering runs). By declaring recessive null alleles for all loci in STRUCTURE, null allele frequencies were estimated and accounted for. The most optimal number of genetic clusters was determined by plotting the log-likelihood of the data Ln *P*(*D*) against the number of clusters *K* (Pritchard et al., 2000) using STRUCTURE HARVESTER (Earl and vonHoldt, 2012), as well as by assessing the stability of replicate runs for each *K* (10 iterations per *K*).

The genetic diversity among the wild and cultivated specimens was summarized using a principal component analysis (PCA) and visualized as a scatterplot with the R (R Development Core Team, 2011) packages *adegenet* (Jombart, 2008) and *ade4* (Chessel et al., 2007).

### Genetic diversity and differentiation

To compare genetic diversity among wild and cultivated *C. canephora* shrubs, as well as among the different sampling locations, the following genetic diversity indices were calculated: number of alleles (*NA*), number of effective alleles (*NAe*), rarefied allelic richness (*AR*), expected heterozygosity (*He*), observed heterozygosity (*Ho*), and inbreeding coefficient (*Fi*). Pairwise genetic differentiation was assessed by calculating *F*_ST_ between wild and cultivated *C. canephora* shrubs, as well as between wild populations from different geographic areas. Furthermore, allele frequencies for the 18 microsatellite loci were calculated and compared among the wild and cultivated specimens to assess the genetic similarity of cultivated and wild *C. canephora* shrubs. All calculations were done using SPAGeDi 1.5d (Hardy and Vekemans, 2002).

## RESULTS

### Genetic population structure and admixture

Four genetic clusters were inferred for our *C. canephora* dataset (Figure 2) using the Bayesian clustering algorithm implemented in STRUCTURE. In this clustering analysis, wild populations were first separated from the cultivated plants sampled in backyards and from the INERA Yangambi collection (at *K* = 2) (Appendix S2). Subsequently, wild populations were subdivided based on geographic origin, separating the specimens collected in Epulu and Djugu (Ituri province) from specimens collected in Yangambi and Yoko (Tshopo province) (at *K* = 3). Lastly, six accessions from the INERA Yangambi collection were separated (at *K* = 4). Out of these six accessions, four were identified as ‘Petit Kwilu’, a variety originating from the Mayombe (western DR Congo and Congo-Brazzaville, Cabinda and coastal Gabon). These six distinct individuals showed increasing levels of admixture at *K* = 5 to *K* = 10 (Appendix S2).

**Figure 2.**
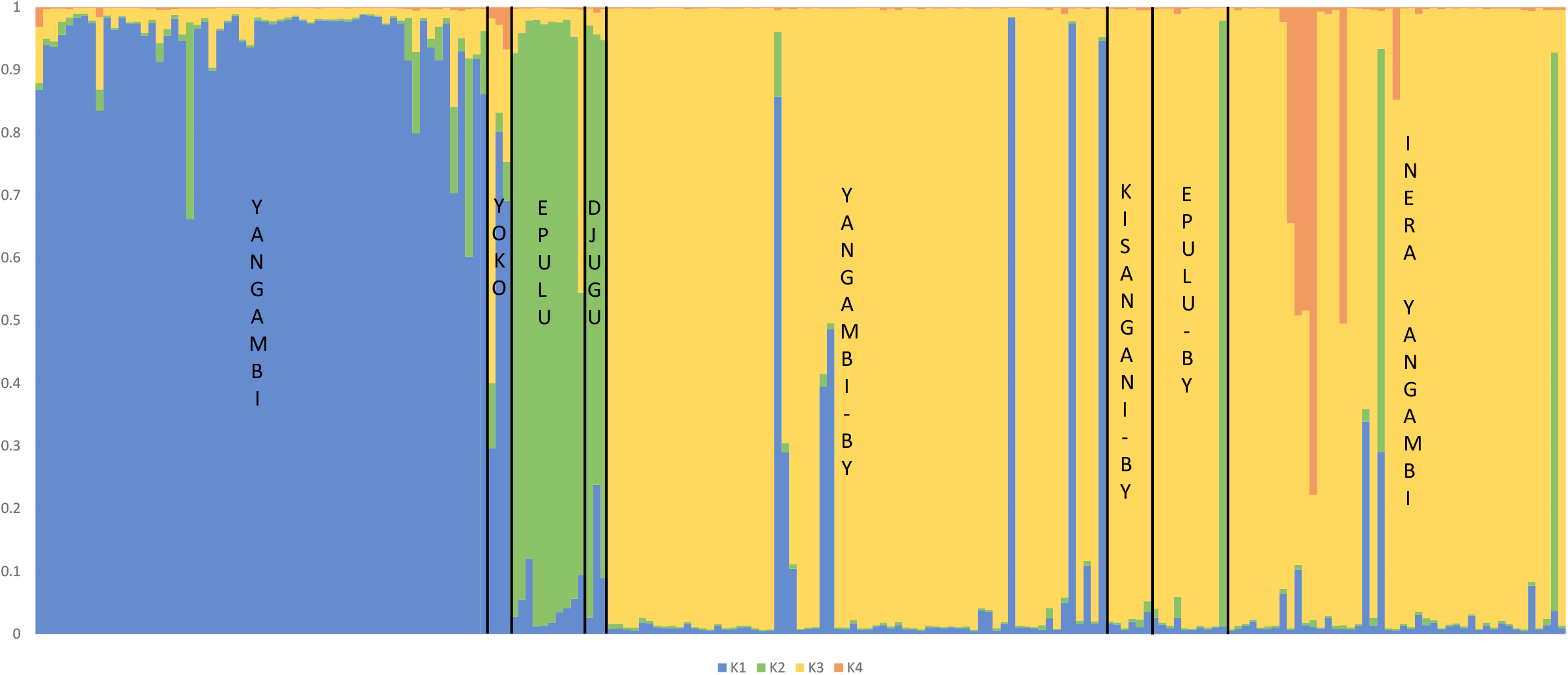
The bar plot representing the assignment probabilities (vertical axis) for the most likely number of genetic clusters *K* = 4 in the *Coffea canephora* microsatellite dataset, inferred using STRUCTURE (Pritchard et al., 2000).

Some of the specimens collected from backyards in Yangambi and Epulu had genotypes that matched the wild genotypes found in the local forest populations (four and one specimen, respectively) (Figure 1, Figure 2). By contrast, no (introduced) wild genotypes were found in the backyards in Kisangani. Overall, six specimens had a ‘hybrid’ wild-cultivated genotype: one in Yoko, one in Epulu, three in Yangambi backyards and one in the INERA Yangambi collection (SY66). One accession (L51Y65) from the INERA Yangambi collection had a genotype that matched the wild genotypes from the Ituri province, and one accession (NA; no accession number available) showed a mix of the wild genotypes from Ituri and Tshopo. The STRUCTURE assignment probabilities also indicated low levels of admixture between wild populations from the Tshopo and Ituri province (Figure 2).

For the STRUCTURE clustering analysis, the plotted log-likelihood of the data Ln *P*(*D*) against the number of clusters *K* showed that the increase in average Ln *P*(*D*) was highest between *K* = 1 and *K* = 2, while the highest average Ln *P*(*D*) was observed at *K* = 4 (Appendix S3). Variability in Ln *P*(*D*) among the 10 iterations was relatively low across all *K*-values.

The PCA (Appendix S4) showed a similar clustering as obtained with STRUCTURE, with the first axis (PC1) mostly separating wild from cultivated specimens. Along the second axis (PC2), the wild populations were separated depending on their geographic origin (Ituri vs. Tshopo province), and the six distinct accessions collected from INERA Yangambi (which were separated at *K* = 4 in the STRUCTURE clustering analysis) were separated from the rest of the cultivated specimens. The third axis (PC3) again separated the wild populations based on their geographic origin, highlighting the relatively high genetic diversity found in the wild populations. The first three principal components explained 6.09%, 3.38% and 2.86% of the variance.

### Genetic diversity and differentiation

Both the number of alleles (*NA*) and the effective number of alleles (*NAe*) were highest for the wild populations (*NA* = 7.94, *NAe* = 3.94), followed by the INERA Yangambi Coffee Collection (*NA* = 7.39, *NAe* = 3.37) and the backyard samples (*NA* = 6.89, *NAe* = 3.09) (Table 1). The allelic richness (*AR*, among 12 gene copies *k*), expected heterozygosity (*He*) and observed heterozygosity (*Ho*) were highest in the INERA Yangambi collection (*AR* = 4.27, *He* = 0.64, *Ho* = 0.53), followed by the wild populations (*AR* = 4.25, *He* = 0.63, *Ho* = 0.47) and the backyard samples (*AR* = 3.83, *He* = 0.59, *Ho* = 0.50). Since a relatively large part of the genetic diversity in the INERA Yangambi collection might originate from the six distinct accessions (LAF159, S23, S19, L6, L251Y128 and NA) that were separated in the PCA and the clustering analysis, genetic diversity indices were also calculated without the respective six accessions. This resulted in lower genetic diversity estimates for the INERA Yangambi collection (*NA* = 6.61, *NAe* = 3.18, *AR* = 4.01, *He* = 0.62 and *Ho* = 0.50), all slightly lower than the genetic diversity measures estimated in the wild populations (except for *Ho*). Among the wild populations, genetic diversity was higher in the Tshopo province (*NA* = 6.83, *NAe* = 3.57, *AR* = 4.04, *He* = 0.61 and *Ho* = 0.48) than in the Ituri province (*NA* = 4.78, *NAe* = 3.45, *AR* = 3.84, *He* = 0.59 and *Ho* = 0.46). Number of alleles (*NA* and *NAe*) might be higher in the Tshopo province because of the larger sampling size, while allelic richness (*AR*) en expected heterozygosity (*He*) are not affected by sampling size. Individual inbreeding coefficients (*Fi*) were significant for all groups and ranged between 0.16 (backyard specimens) and 0.26 (wild populations) (Table 1).

**Table 1.** Genetic diversity parameters for wild and cultivated *Coffea canephora* populations.

| Population | *n* | *NA* | *NAe* | *AR* (k=12) | *He* | *Ho* | *Fi* |
| --- | --- | --- | --- | --- | --- | --- | --- |
| Wild | 69 | 7.94 | 3.94 | 4.25 | 0.64 | 0.47 | 0.26 |
| *Tshopo province* | 56 | 6.83 | 3.57 | 4.04 | 0.61 | 0.48 | 0.22 |
| *Ituri province* | 13 | 4.78 | 3.45 | 3.84 | 0.59 | 0.46 | 0.24 |
| Backyards | 81 | 6.89 | 3.09 | 3.83 | 0.59 | 0.50 | 0.16 |
| *Backyards -5ind* | 76 | 6.33 | 2.98 | 3.68 | 0.58 | 0.50 | 0.13 |
| INERA Yangambi Collection | 45 | 7.39 | 3.37 | 4.27 | 0.64 | 0.53 | 0.18 |
| *Collection -6ind* | 39 | 6.61 | 3.18 | 4.01 | 0.62 | 0.50 | 0.20 |
| *All* | 195 | 10.67 | 4.03 | 4.57 | 0.67 | 0.50 | 0.26 |

Notes: *n* number of individuals analyzed, *NA* number of alleles, *NAe* effective number of alleles (Nielsen et al., 2003), *AR* (k = x) allelic richness or number of alleles among x gene copies, *He* expected heterozygosity corrected for sample size, *Ho* observed heterozygosity, *Fi* individual inbreeding coefficient. *Fi* was significantly larger than 0 for all populations. “Backyards -5ind” are all the cultivated backyard individuals without the five individuals with a wild genotype. “Collection -6ind” are all the cultivated individuals from the INERA Yangambi Coffee Collection without the six genetically distinct individuals (mostly ‘Petit Kwilu’ accessions).

The allele frequencies calculated for the 18 microsatellite loci in the wild and cultivated specimens showed that all loci harbored alleles that were unique to at least one of both categories (Table 2): three loci harbored alleles that were unique to cultivated specimens (*R325, SSR209, R342*), while the other 15 loci harbored unique alleles for both cultivated and wild specimens. For one locus (*SSR196*, 14 alleles), only unique alleles were observed, either to wild or to cultivated specimens. Among all loci, we found 52 alleles which were unique to wild specimens and 48 alleles unique to cultivated specimens. Nine of those 48 alleles were only present in the six cultivated specimens with a distinct genotype, collected from INERA Yangambi collection. Furthermore, we found five alleles among all loci which were only present in wild specimens and in those six distinct cultivated specimens, but not in any of the other cultivated specimens.

**Table 2.** The number (#) of alleles per locus that were unique to wild or cultivated specimens. The number between brackets () indicates the number of alleles that were unique to the six genetically distinct individuals from the INERA Yangambi Coffee Collection (mostly ‘Petit Kwilu’ accessions).

| Locus | # alleles | # alleles unique to<br>wild specimens | n° alleles unique to<br>cultivated specimens |
| --- | --- | --- | --- |
| R338 | 13 | 2 | 4 (1) |
| SSR146 | 9 | 2 | 3 (1) |
| R278 | 11 | 5 | 1 |
| R339 | 17 | 6 | 1 |
| R336 | 14 | 2 | 4 |
| R325 | 5 | 0 | 3 |
| SSR196 | 14 | 8 | 6 (1) |
| R268 | 7 | 3 | 2 (1) |
| R168 | 11 | 3 | 1 |
| R189 | 9 | 1 | 2 |
| SSR497 | 19 | 7 | 1 |
| R175 | 7 | 2 | 1 |
| SSR495 | 8 | 2 | 3 (1) |
| R250 | 15 | 2 | 6 |
| R148 | 9 | 4 | 1 |
| SSR209 | 8 | 0 | 4 (3) |
| R342 | 4 | 0 | 1 |
| SSR533 | 11 | 3 | 4 (1) |
| *Total* | 191 | 52 | 48 |

Pairwise genetic differentiation (*F*_ST_) (Table 3) was highest between the wild populations and the specimens collected from backyards (*F*_ST_ = 0.144) or from the INERA Yangambi collection (*F*_ST_ = 0.135). Genetic differentiation was low between the specimens collected from backyards and from the INERA Yangambi collection (*F*_ST_ = 0.004). Pairwise genetic differentiation between the wild populations from the Tshopo and Ituri provinces was slightly lower than the differentiation between all wild populations and cultivated accessions in backyards and the INERA Yangambi Coffee collection (*F*_ST_ = 0.119).

**Table 3.** Genetic differentiation (*F*_ST_) between wild and cultivated specimens of *Coffea canephora*, and among wild populations from the Tshopo and Ituri provinces in northeastern DR Congo.

| $F_{ST}$ | Wild | Backyard | $F_{ST}$ | Tshopo |
| --- | --- | --- | --- | --- |
| **Wild** |  |  | **Tshopo** |  |
| **Backyards** | 0.144 |  | **Ituri** | 0.119 |
| **INERA Collection** | 0.135 | 0.004 |  |  |

## DISCUSSION

The distribution of the genetic diversity of wild and cultivated *C. canephora* in the Tshopo and Ituri provinces revealed some unexpected patterns. Firstly, the agreement in the genetic constitution of the INERA Yangambi accessions and the vast majority of the shrubs in the backyard gardens, indicates that most people growing *C. canephora* locally, received their material directly or indirectly from INERA breeding programs. Secondly, the cultivated shrubs, both from backyards and the INERA collection, are genetically clearly distinct from the local wild gene pool, showing relatively large genetic differentiation. Levels of genetic diversity are similar for the INERA and the wild populations, but both wild and cultivated specimens harbor a lot of unique alleles. Finally, the wild gene pool from the Tshopo and Ituri provinces is not represented in the INERA Yangambi collection. These observations have important conservation implications as will be discussed below.

### Genetic diversity, structure and origin of wild Coffea canephora

Genetic diversity of *C. canephora* was slightly higher in the Tshopo province as compared to the shrubs sampled in Ituri, although expected and observed heterozygosity were fairly similar. This can be explained by the broader geographic sampling in the Tshopo province, which included wild populations from both the Yangambi and Kisangani region (incl. Yoko). The expected heterozygosity in the Tshopo and Ituri provinces (*He* ∼ 0.60) matched that of the most diverse populations in Uganda (Kiwuka et al., 2021) and that of some of the most diverse undisturbed *C. arabica* stands (Aerts et al., 2013). Both studies used SSR markers with respectively 19 and 24 microsatellite loci, as compared to 18 in our study. The values of expected heterozygosity are at the upper end of those estimated for diversity of an outcrossing perennial plant using microsatellite markers (0.47 to 0.68; Nybom, 2004). The pronounced self-incompatibility system in *C. canephora* (Lashermes et al., 1996) seems to ensure the maintenance of high levels of heterozygosity within populations.

Relatively strong genetic differentiation (*F*_ST_ = 0.119) was observed between wild populations from the Tshopo province and Epulu, which are separated about 400 km from each other. A more pronounced genetic structure is found amongst wild *C. canephora* populations in Uganda, which can be explained by the more pronounced (historic and/or present-day) population fragmentation in the area (Kiwuka et al., 2021). The tropical rainforest in the Congo Basin is still unfragmented and it can be expected that *C. canephora* shrubs are distributed somewhat continuously throughout this rainforest. However, past glaciations (e.g. during the Pleistocene) drastically reduced the rainforest cover (Maley, 1996; Anhuf, 2000; Gomez et al., 2009; Hardy et al., 2013), which caused genetic differentiation between populations in isolated forest refugia due to genetic drift, bottlenecks and inbreeding. Such past barriers to gene flow could (partly) explain the observed differentiation between the populations in Tshopo and Ituri, possibly in combination with present-day dispersal barriers. While little is known about pollinator specificity in wild *C. canephora* shrubs and the distances the pollinators can cover, it has been suggested that long distance seed dispersal of the red *Coffea* berries by birds and perhaps mammals could potentially reach up to 100 kilometers, thus contributing significantly to gene flow across large distances (Charrier, 1971; Berthaud, 1986). However, this claim of long-distance dispersal has been disputed due to the fact that birds living in the rainforest understory commonly have a sedentary habit and a rapid gut passage (Theim et al., 2014; Grant et al., 2019).

### Diversity and origin of the INERA Yangambi collection

The majority of the accessions in the INERA Yangambi collection are referred to as ‘Lula’ varieties (Table S1) and presumably originate from the Lula research station near Kisangani. The wild origin of the Lula variety remains unclear, but given the high level of distinct alleles in the wild and cultivated shrubs it seems unlikely that the origin is to be found in our sampling region, i.e. Tshopo and Ituri. Nonetheless, the ‘Lula’ varieties are assumed to have originated from the Congo Basin and probably root back to the early introduction of ‘*Coffea robusta’* by Linden from Sankuru, but additional sampling and research is needed to trace the region of origin. Cultivated material from the former INEAC Yangambi collection is still present in the CNRA collection in Ivory Coast (Cubry et al., 2013), but this cultivated material dates back to 1935 (Bodard, 1965). The CNRA *C. canephora* material originating from the former INEAC was shown to be closely related to *C. canephora* growing in Uganda (Cubry et al., 2013; Leroy et al., 2014). However, further studies are required to assess the relationships between the ‘Lula’ variety currently present in the INERA Yangambi Coffee Collection and the material distributed to CNRA during the INEAC period and the Ugandan gene pool. The relatively high genetic variation in the INERA Yangambi Coffee Collection, as expressed by the high levels of heterozygosity and allelic richness, can be explained by the diverse origin of several rare genetic lines in the collection. Four ‘Petit Kwilu’ accessions, originating from the Mayombe Region (western DR Congo, Congo and Gabon), in the collection were genetically clearly distinct from the ‘Lula’ varieties, as was confirmed by previous studies (e.g. Leroy et al., 2014). In addition, accessions originating from the North Kivu, Haute Zaire and Equateur provinces, with one representative each, were also present in the collection (Table S1), although it must be noted that provenances are not well documented in the collection. Very little (local) wild genetic diversity seems to be preserved in the INERA Yangambi Coffee Collection. The introduction of the local wild genetic diversity into the INERA field collection is a relatively easy, but very important, way to enrich Robusta coffee genetic resources in the collection. The availability of local genetic resources in the collection could potentially be a useful source for breeding of *C. canephora* shrubs that are adapted to local soil and climatic conditions. In addition, ex situ conservation of local genetic resources, which are threatened by deforestation, change in forest structure and the disappearance of seed dispersers (Sellan et al., 2017; van Vliet et al., 2018; Kyale Koy et al., 2019), can complement in situ conservation efforts.

### Backyard garden cultivation

Genetic differentiation between *C. canephora* shrubs in the INERA Yangambi Coffee Collection and the shrubs in backyards was very low. The vast majority of the coffee plants in backyards are most likely ‘Lula’ varieties originating from the INERA breeding program, that have been distributed to local villagers. Even the cultivated plants in the Ituri province were closely related to the ‘Lula’ variety. Since coffee grown in backyards in the study region is used mainly for own consumption and to make a decoction from the leaves (Campa et al., 2012), the dominance of non-local ‘Lula’ varieties was somewhat surprising. The lower genetic diversity of backyard shrubs can be explained by the fact that mainly seven ‘Lula’ elite lines, originating from the INERA Yangambi collection, are used for germplasm production and distribution (Tshimi Ebele, pers. comm.). Since breeding activities at the INERA Yangambi have reduced drastically over the last decade, it can be expected that the genetic resources that have been distributed have remained relatively uniform. The large gene pool of wild *C. canephora* shrubs in the Tshopo and Ituri provinces was very poorly represented in the backyard cultivation systems of local villagers, despite the fact that they are sometimes separated by a kilometer or less. Moreover, genetic differentiation was highest between the wild and the backyard gene pool, and only 5 out of 81 samples (6.2%) collected in backyards could be assigned a local wild origin. This observation contrasts somewhat with the situation for *C. canephora* in Uganda, where gene pools of cultivated and wild plants are much more mixed (Musoli et al., 2009; Kiwuka et al., 2021). This lower genetic differentiation between cultivated and wild gene pools in Uganda is likely due to a more extensive use of local wild genetic resources during Robusta cultivation, as well as a more recent and less extensive selection process.

Due to the close vicinity of cultivated *C. canephora* shrubs (sometimes less than 1 km) and the limited domestication (wild shrubs are morphologically quite similar to cultivated material), a considerable impact of cultivated shrubs on the integrity of the wild gene pool could be expected. Indications for crop-wild introgression in coffee were observed in a study of *C. arabica* populations in Ethiopia (Aerts et al., 2013) and *Coffea canephora* in Uganda (Kiwuka et al., 2021), yet we found little evidence for gene flow from the cultivated gene pool into the wild gene pool in the NE Congo Basin. Any attempt to explain this contrasting observation would be speculative at this point, and a broader sampling is needed to confirm this observation. Five putative crosses between wild local shrubs and cultivated *C. canephora* shrubs were, however, observed in backyards of Yangambi, Yoko and Epulu. The origin of such ‘hybrids’ appears to be unclear, yet the most reasonable explanation would be a crossing event from wild and cultivated *C. canephora* growing together in backyards.

## CONCLUSIONS

The present findings show that the cultivated *Coffea canephora* accessions from INERA Yangambi and the vast majority of the shrubs in the backyard gardens in northeastern DR Congo are genetically very similar. This indicates that most people growing *C. canephora* locally received their material directly or indirectly from INERA breeding programs, while a few shrubs are obtained directly through collections from surrounding forests. Furthermore, the cultivated shrubs, both from backyards and the INERA collection, are genetically distinct from the local wild gene pool, showing relatively large genetic differentiation and both gene pools harbor multiple unique alleles. The introduction of the local wild genetic diversity into the INERA field collection would be a great way to increase the genetic resources available for breeding purposes, as well as to support the ex situ conservation of *C. canephora*. The application of high-throughput sequencing methods in future studies would be beneficial to characterize putative introgression and gene flow between cultivated shrubs and local wild populations, often growing in close proximity. Such genetic studies could be complemented with studies focused on pollen and seed dispersers, since little is known about the gene dispersal mechanisms in *C. canephora*, and in tropical understory shrubs in general.

## ACKNOWLEDGEMENTS

The authors want to thank Wim Baert (Meise Botanic Garden) and Pieter Asselman (UGent) for support with molecular lab work. We are grateful to the Ministère de L’Environnement et Développement Durable and Institut National pour l’Étude et la Recherche Agronomiques (INERA), Democratic Republic of Congo for their help with obtaining collecting permits. This study was financially supported by EU XI^th^ Development Fund (FORETS project), the Foundation for the promotion of biodiversity research in Africa (SBBOA, www.sbboa.be), Research Foundation - Flanders (Research project G090719N), the Belgian Science Policy (BELSPO; grant B2/191/P1/COFFEEBRIDGE). SVA received funding from the Belgian American Educational Foundation (www.baef.be), JD received funding from the Research Foundation - Flanders (FWO; 1125221N).

## AUTHOR CONTRIBUTIONS

FV, SVA, PS and SBJ conceived and designed the study. SVA, FV, PS and SBJ wrote the manuscript. YB did the molecular lab work. SVA did the data analysis. FV, SN, JAA, BK, IMM, SVA and PS collected data and samples. All authors read and approved the final version of the manuscript.

## DATA AVAILABILITY

The data generated in this study is available from the authors upon reasonable request.

## SUPPORTING INFORMATION

Additional supporting information may be found online in the Supporting Information section at the end of the article

APPENDIX S1. Sequences for the microsatellite primers used in the present study, including fluorescent Q-tail and multiplex information.

APPENDIX S2. The bar plots for *K* = 2 to *K* = 4, and *K* =10 representing the assignment probabilities (y-axis) inferred using STRUCTURE, for the complete *Coffea canephora* microsatellite dataset.

APPENDIX S3. Likelihood of the *Coffea canephora* microsatellite dataset as a function of the assumed number of genetic clusters (*K*) according to the Bayesian clustering algorithm implemented in STRUCTURE. The circle represents the mean log-likelihood over 10 runs, while the vertical bars show the standard deviation among runs.

APPENDIX S4. Principal Component Analysis (PCA) of the genetic diversity in the *Coffea canephora* microsatellite dataset. *PC* stands for Principal Component.

## APPENDICES

Appendix 1. List of wild and cultivated *Coffea canephora* accessions from northeastern Democratic Republic of the Congo included in the current study.

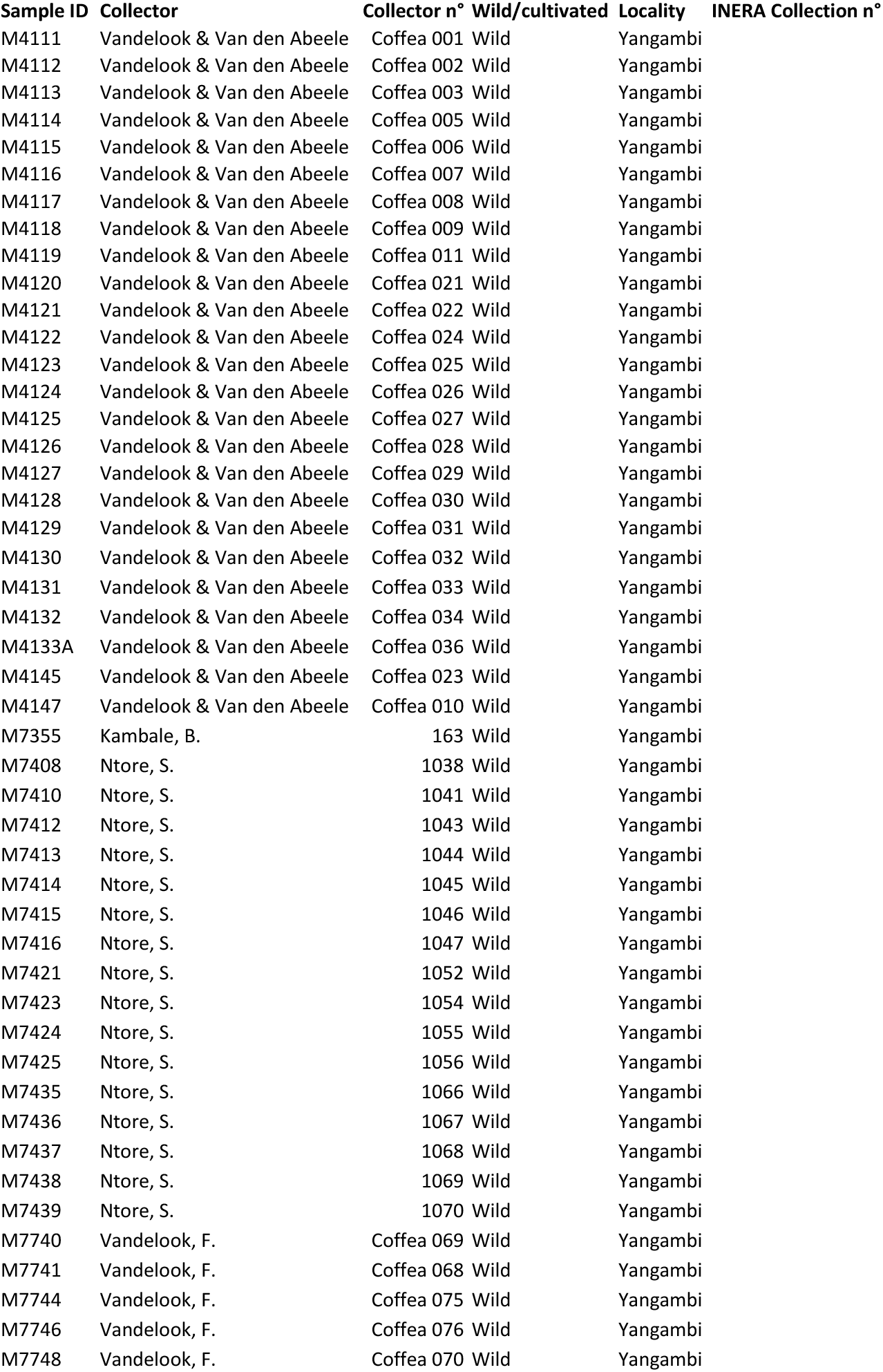
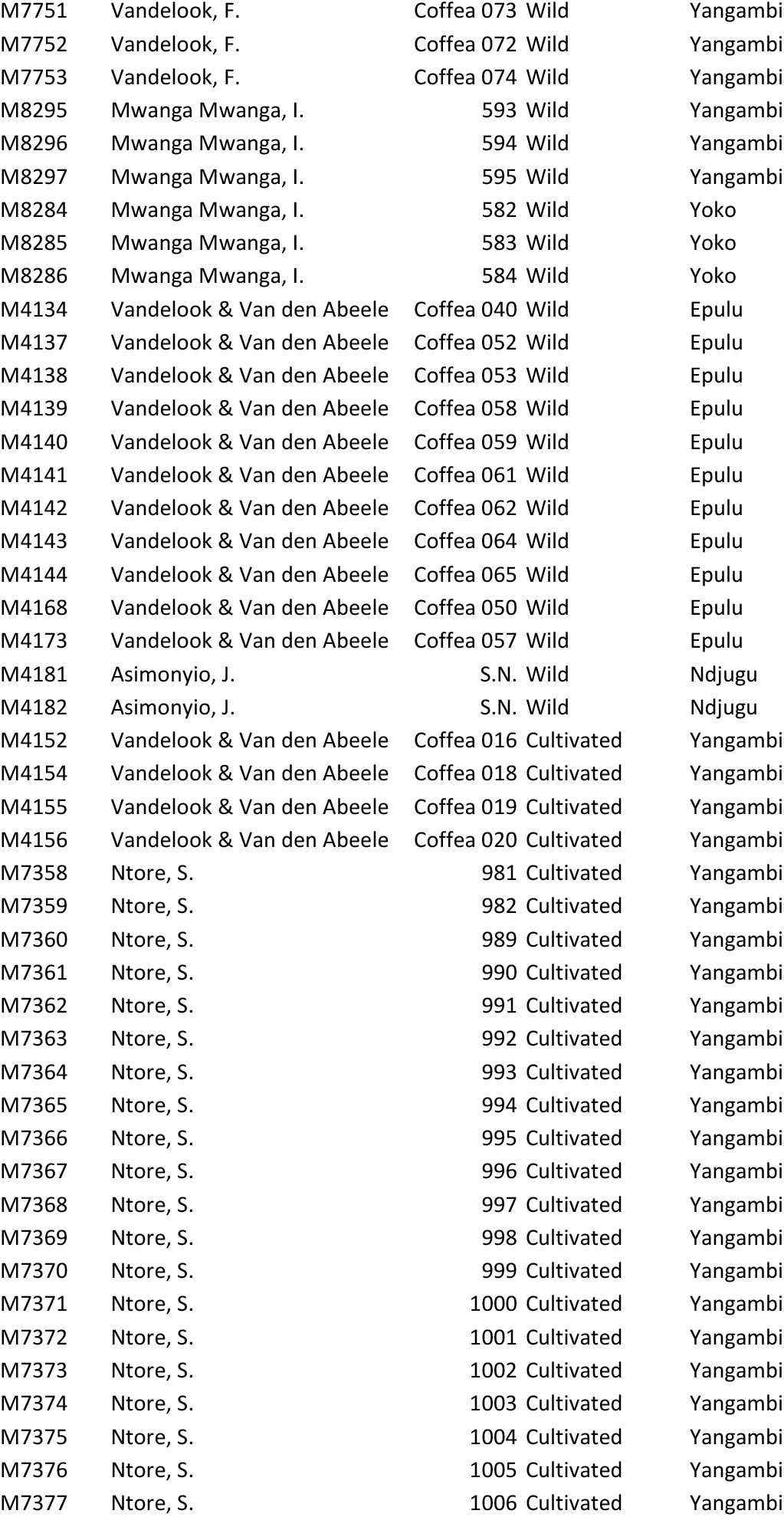
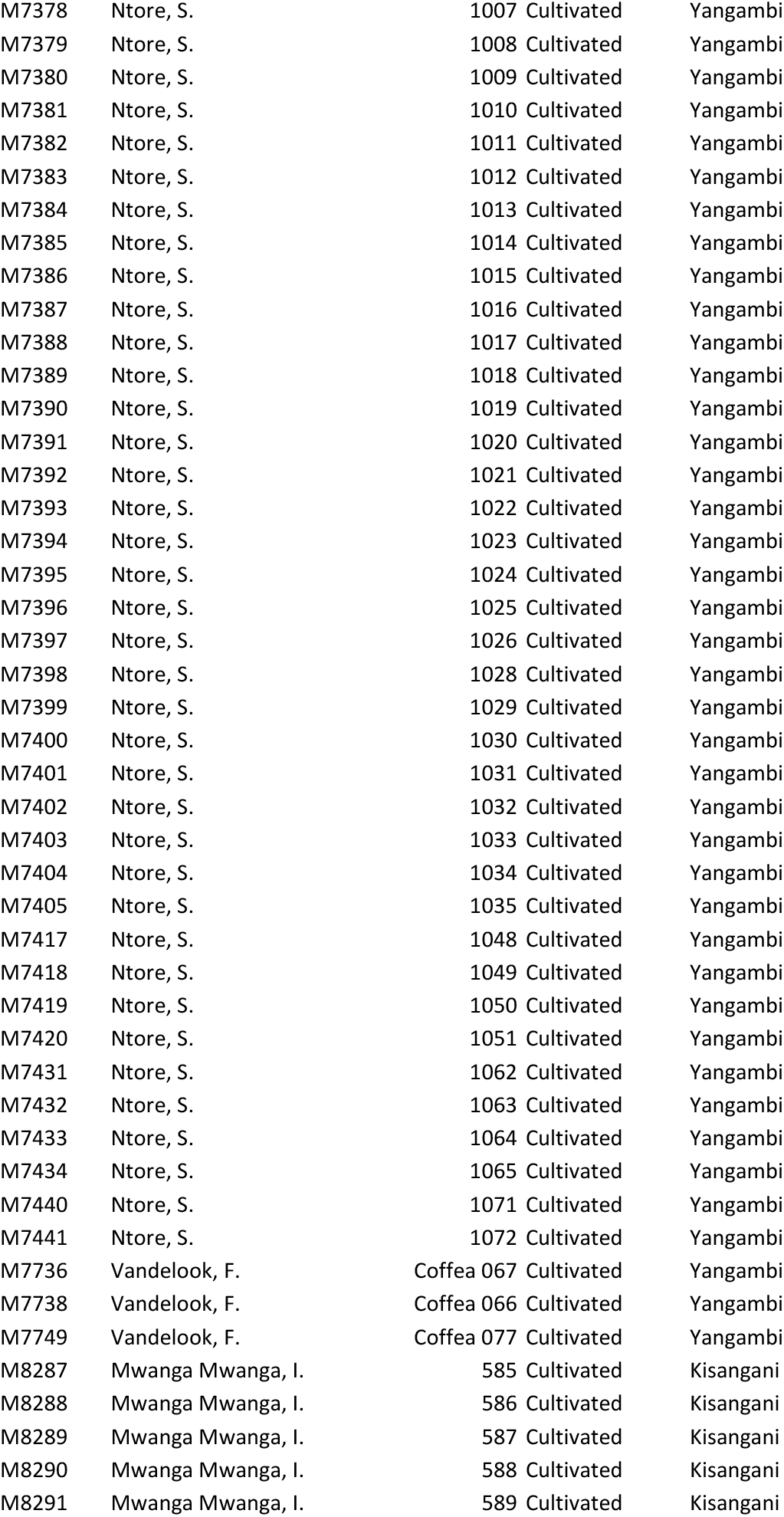
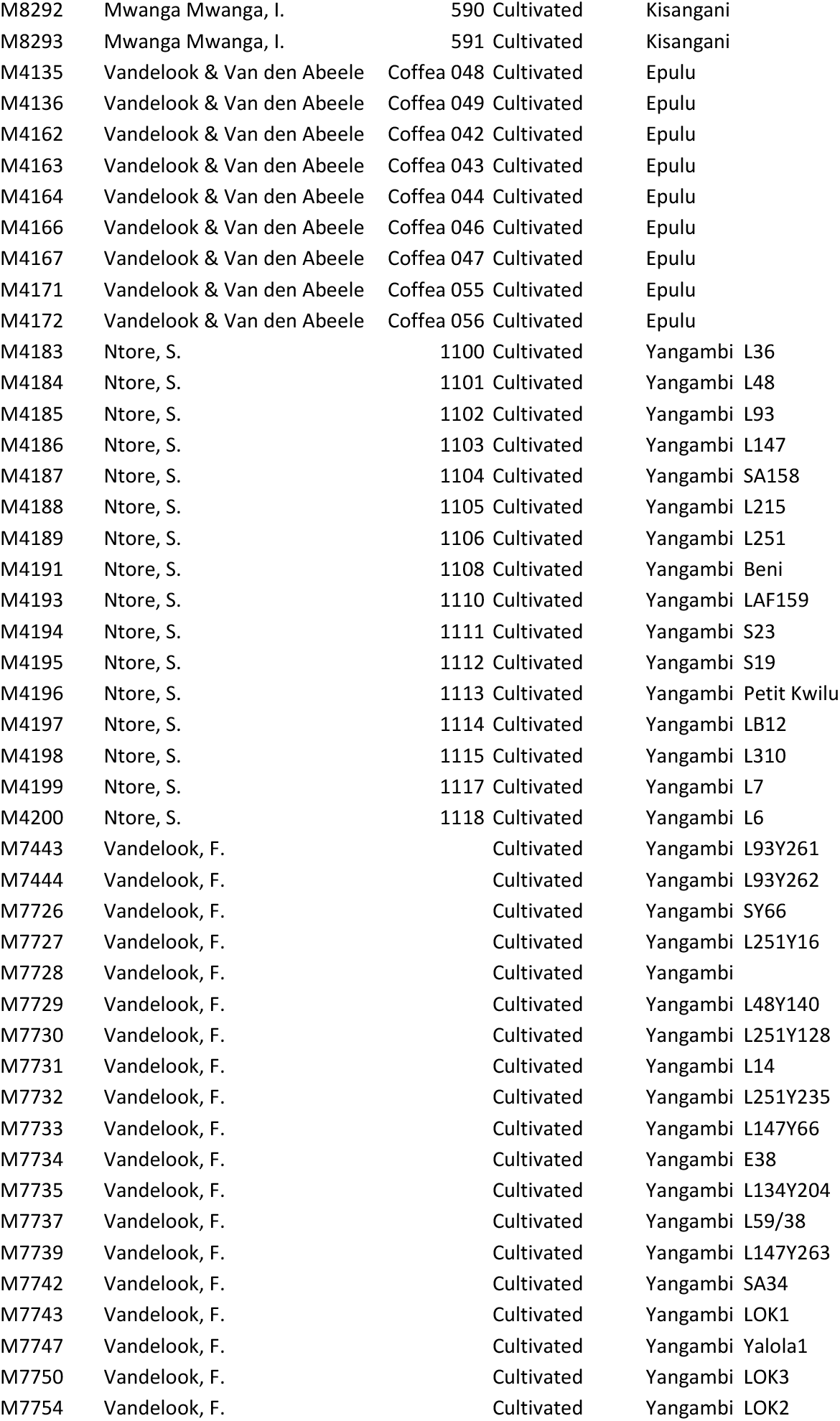
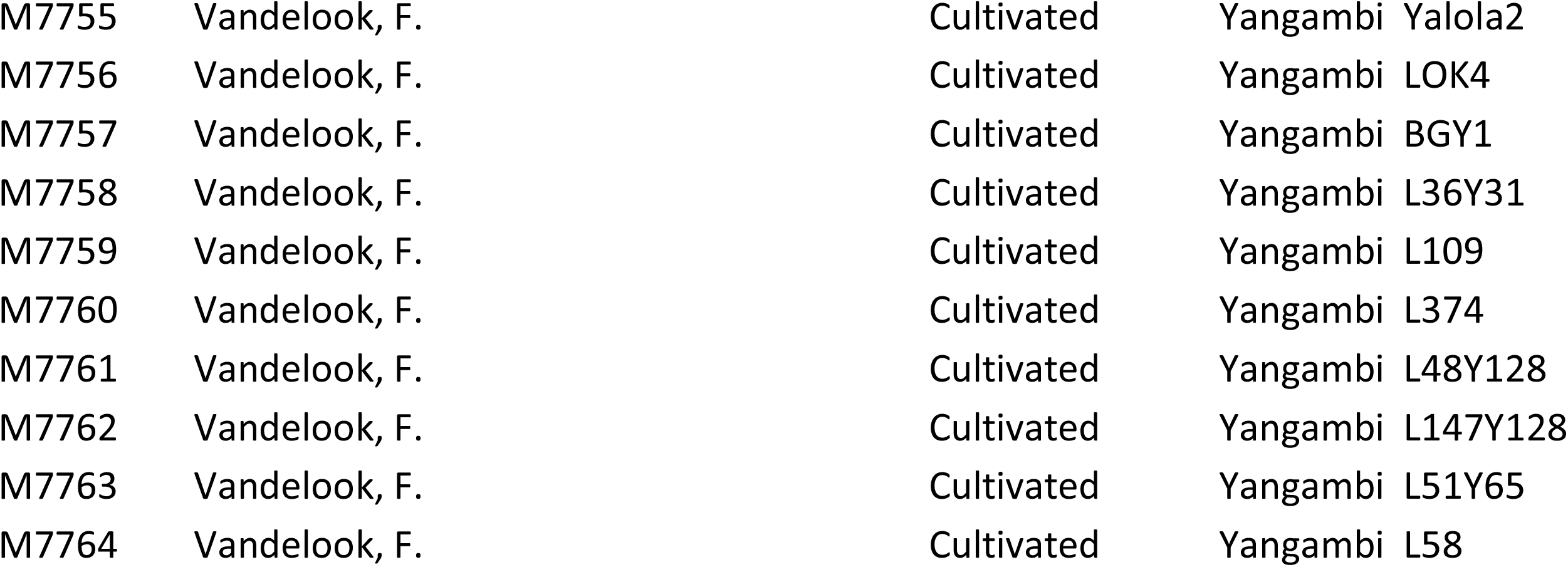

