## Appendix S1 for "Genetic diversity of wild and cultivated *Coffea* canephora in northeastern DR Congo and the implications for conservation"

| *Locus* | *Primer sequences (5'-3')* | *Labelled primer* |
|---|---|---|
| **Multiplex 1** | | |
| R338 | F: CGAAGGCTGTCAACAACTGG | Q1-6-FAM |
| | R: TGTAAAACGACGGCCAGTGGGATAAACAAGTTAAAGGA | |
| SSR146 | F: TCCAGTTTGATCAGCAACCA | Q1-6-FAM |
| | R: TGTAAAACGACGGCCAGTCCATCTTGGGGATAGAGCAA | |
| R278 | F: TGTAGATTTGAAACCCAATC | Q3-VIC |
| | R: CACTGCTTAGAGCGATGCAAGTCTCGACAAGTTTTGAC | |
| R339 | F: ATTATGCTCGCTGGGCTGTT | Q3-VIC |
| | R: CACTGCTTAGAGCGATGCTGGGATCACTCCTGTGTCGC | |
| R336 | F: TTGCCTTTTTAGTGCGTGTA | Q4-PET |
| | R: CTAGTTATTGCTCAGCGGTGCAAAGCCCGAGGATT | |
| **Multiplex 2** | | |
| R325 | F: CCTTGTTGTTGGGGAATGTC | Q1-6-FAM |
| | R: TGTAAAACGACGGCCAGTGGCTGTTCTGGGCTTTGTG | |
| SSR196 | F: ATCCCCATCAGAAGACCTCA | Q1-6-FAM |
| | R: TGTAAAACGACGGCCAGTCCTCCACCGCCTGTTTATTA | |
| R268 | F: GTATCCCACAATGAAATCAC | Q2-NED |
| | R: TAGGAGTGCAGCAAGCATAGTAGAATTTTCAACATATAAG | |
| R168 | F: CCTGGACTGGTAGAAACAAA | Q4-PET |
| | R: CTAGTTATTGCTCAGCGGTAAAGGTGTTCAATGCCTACA | |
| **Multiplex 3** | | |
| R189 | F: GGAGTGAGAGGAGGGCGTAG | Q1-6-FAM |
| | R: TGTAAAACGACGGCCAGTGAGAGAGGGACACTGCTGC | |
| SSR495 | F: TCGGCTCCCAAATATTCATC | Q4-PET |
| | R: CTAGTTATTGCTCAGCGGTCATGAGGCAAGAGGGTTTGT | |
| SSR497 | F: AGGCTTGCTGGAACTCTTGA | Q3-VIC |
| | R: CACTGCTTAGAGCGATGCGAAAGACTTGTCCTTTGCCG | |
| R175 | F: GCAGTGACGCAGCAATG | Q2-NED |
| | R: TAGGAGTGCAGCAAGCATAAAAGGAGAGCCAAAGCAGT | |
| **Multiplex 4** | | |
| R250 | F: GCATCATTGGGTTGGTGG | Q1-6-FAM |
| | R: TGTAAAACGACGGCCAGTCGACTTTCCGCACGCAAAC | |
| R148 | F: CGTCGTTCGAGGACTTGTTC | Q3-VIC |
| | R: CACTGCTTAGAGCGATGCTTCGCAATCCCAGACCC | |
| SSR209 | F: GCCGTGGTGGAAGATGTACT | Q3-VIC |
| | R: CACTGCTTAGAGCGATGCCGAGTTCACCAAGAACGTCA | |
| R342 | F: GCGAGAATAAGGAGTGACC | Q2-NED |
| | R: TAGGAGTGCAGCAAGCATGTCCCTTTTTGTCTGGACC | |
| SSR533 | F: ATCTCCTCGTTCTTCCCCAT | Q4-PET |
| | R: CTAGTTATTGCTCAGCGGTGCTTGTAGCAGGCAGGAAAC | |
