## Supplementary figures and images for "Genetic diversity of wild and cultivated *Coffea* canephora in northeastern DR Congo and the implications for conservation"

### Appendix S2

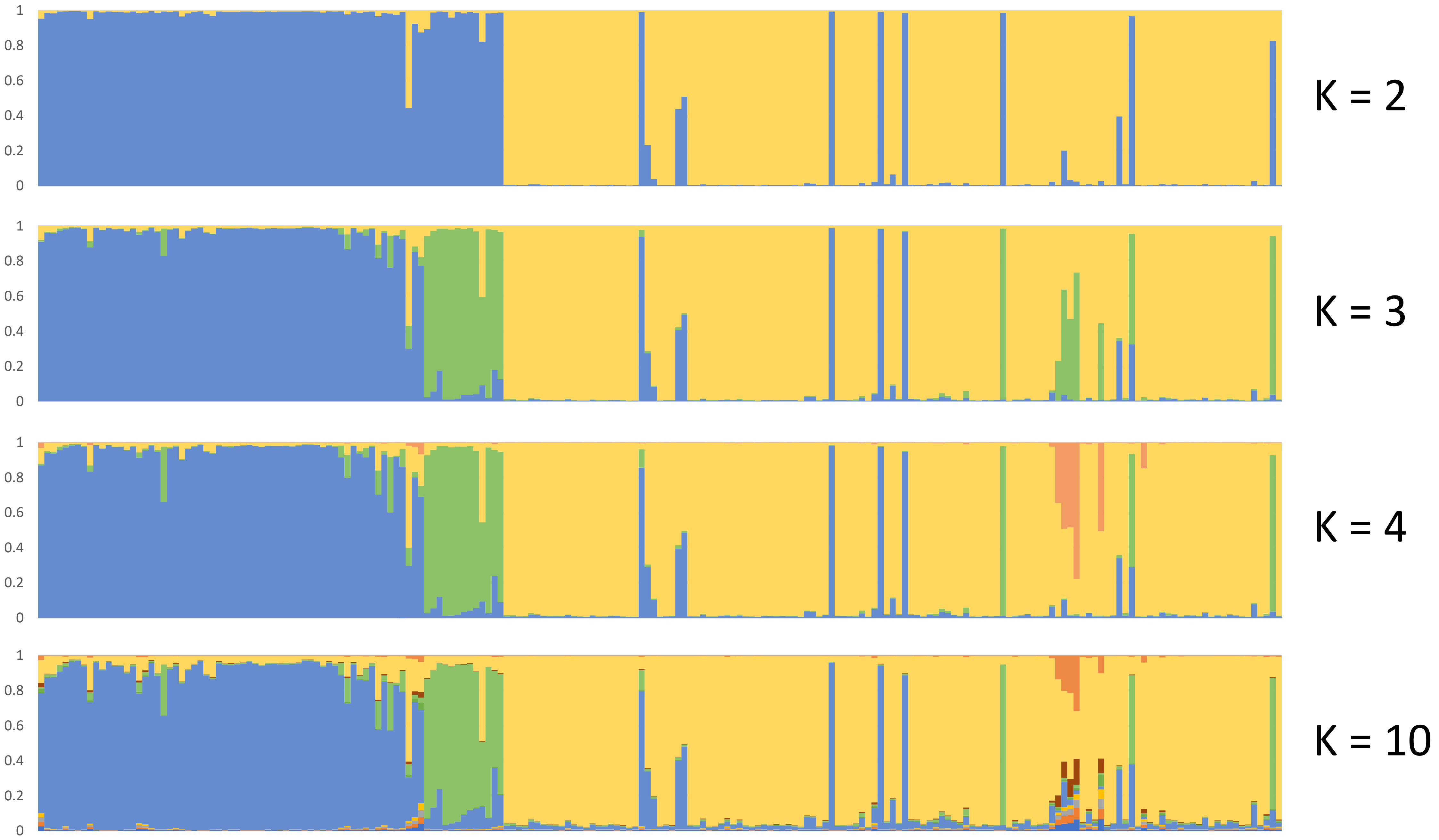

### Appendix S3

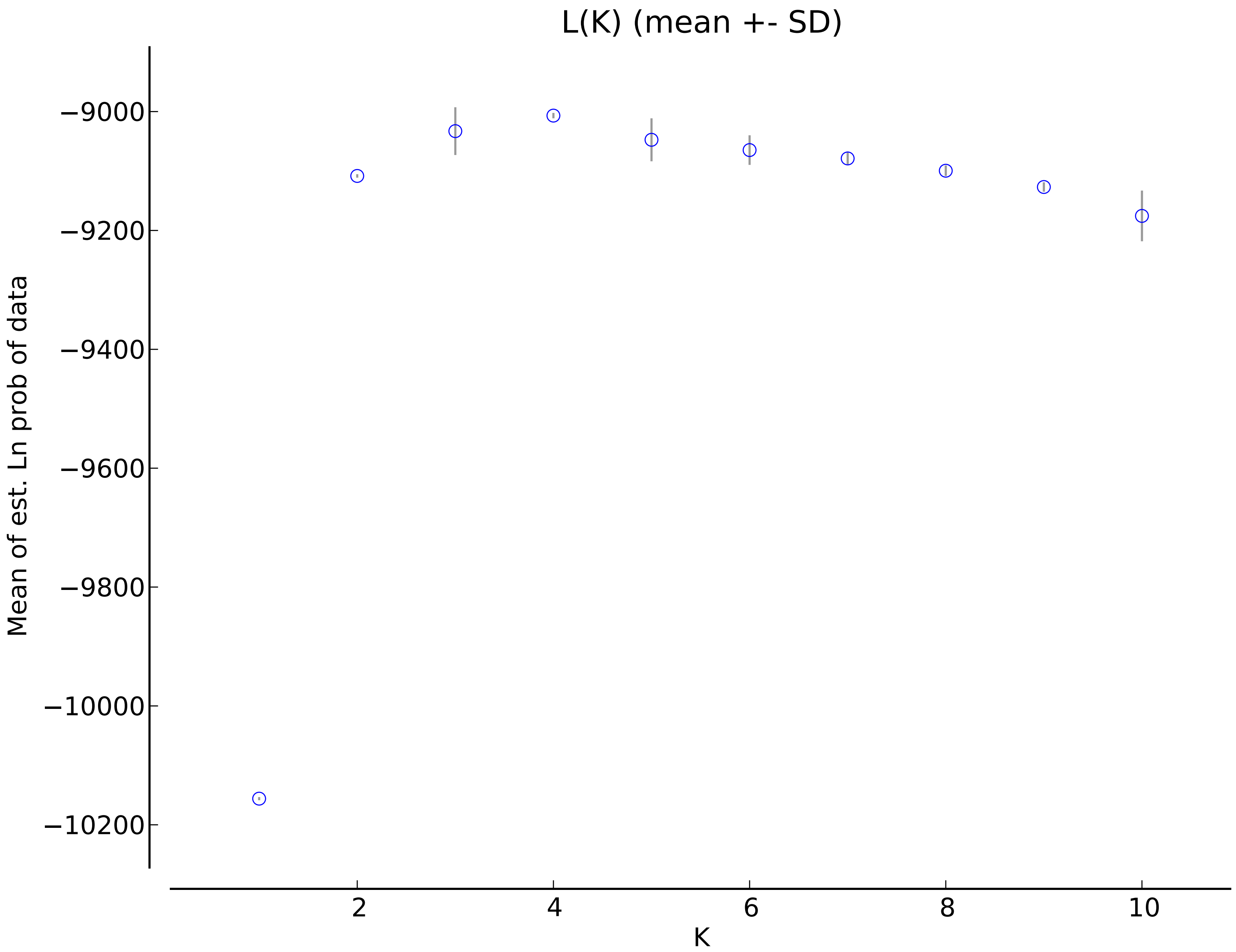
