## Appendix S4 for "Genetic diversity of wild and cultivated *Coffea* canephora in northeastern DR Congo and the implications for conservation"

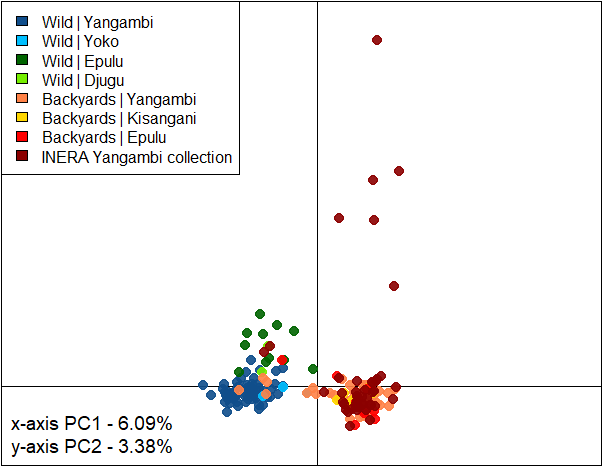

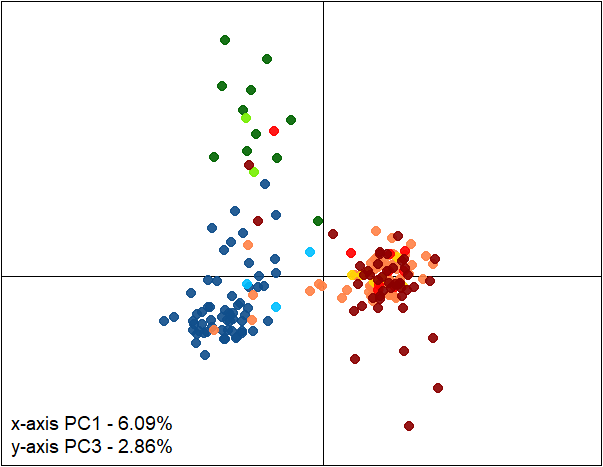

APPENDIX S4. Principal Component Analysis (PCA) of the genetic diversity in the *Coffea canephora* microsatellite dataset. *PC* stands for Principal Component.
